# Real-time visualization of phagosomal pH manipulation by *Cryptococcus neoformans* in an immune signal-dependent way

**DOI:** 10.1101/2022.09.08.507177

**Authors:** Emmanuel J. Santiago-Burgos, Peter V. Stuckey, Felipe H. Santiago-Tirado

**Author notes:** **Correspondence:** Felipe H. Santiago-Tirado. These authors contributed equally to this work (Emmanuel J. Santiago-Burgos, Peter V. Stuckey).

## Abstract

Understanding of how intracellular pathogens survive in their host cells is important to improve management of their diseases. This has been fruitful for intracellular bacteria but it is an understudied area in fungal pathogens. Here we start elucidating and characterizing the strategies used by one of the commonest fungal pathogens, *Cryptococcus neoformans*, to survive intracellularly. The ability of the fungus to survive inside host cells is one of the main drivers of disease progression, yet it is unclear whether *C. neoformans* resides in a fully acidified, partially acidic, or neutral phagosome. Using a dye that only fluoresce under acidic conditions to stain *C. neoformans*, a hypha-defective *Candida albicans* mutant, and the nonpathogenic *Saccharomyces cerevisiae*, we characterized the fungal behaviors in infected macrophages by live microscopy. The main behavior in the *C. albicans* mutant strain and *S. cerevisiae*-phagosomes was rapid acidification after internalization, which remained for the duration of the imaging. In contrast, a significant number of *C. neoformans*-phagosomes exhibited alternative behaviors distinct from the normal phagosomal maturation: some phagosomes acidified with subsequent loss of acidification, and other phagosomes never acidified. Moreover, the frequency of these behaviors was affected by the immune status of the host cell. We applied the same technique to a flow cytometry analysis and found that a substantial percentage of *C. neoformans*-phagosomes showed impaired acidification, whereas almost 100% of the *S. cerevisiae*-phagosomes acidify. Lastly, using a membrane-damage reporter, we show phagosome permeabilization correlates with acidification alterations, but it is not the only strategy that *C. neoformans* uses to manipulate phagosomal acidification. The different behaviors described here provide an explanation to the confounding literature regarding cryptococcal-phagosome acidification and the methods can be applied to study other intracellular fungal pathogens.

## 1 Introduction

*Cryptococcus neoformans* is a major human pathogen with worldwide distribution (1). Because it is found ubiquitously in the environment, we frequently inhale it, establishing a lung infection that in the immunocompetent individual is controlled, but in the immunocompromised it disseminates with tropism for the central nervous system, causing lethal meningoencephalitis. This infection is one of the original AIDS-defining illnesses and still is a leading cause of death among HIV patients, responsible for close to 200,000 deaths yearly in this population (2, 3). Since it is acquired by inhalation, the first line of defense are the alveolar macrophages and other lung phagocytes, which are able to recognize and phagocytose the fungus, but cannot efficiently kill it. The interactions between these host cells and the fungi are critical for the disease and will influence events such as clearance, latency, and dissemination (4). One of the most important interactions driving disease establishment is *Cryptococcus*’s ability to survive inside immune cells such as macrophages.

Normally, when microbes are phagocytosed by macrophages, they end up inside a membranous compartment known as a phagosome. This phagosome follows a well-studied process of maturation that results in the formation of a phagolysosome for destruction of the microbe. A property of a fully functional phagolysosome is its low pH (4.0 – 4.5), conditions that not only are inhospitable to many organisms, but also result in activation of many proteases and other antimicrobial responses. Intracellular pathogens have evolved fascinating strategies to evade this process and survive in cells designed to eradicate them, and one of the most common strategies involve inhibition of phagosomal maturation (5). Some pathogens, such as *Listeria* or *Mycobacteria*, escape from the phagosome before it acidifies; others, such as *Salmonella* or *Legionella*, remain in a modified phagosome that does not fuse with lysosomes. *C. neoformans* is known to reside in a phagosome, but it is unclear if this compartment is a fully-functional phagolysosome or an altered phagosome, and if this compartment acidifies normally is controversial. Some studies suggest it is an acidified lysosome and that acidification is important for fungal replication (6-8). Other results suggest it never, or partially, acidifies (9-12). Moreover, others have seen indirect evidence that *C. neoformans* damages the phagosomal membrane, which would affect its acidification because contents would be neutralized by the cytoplasm (13-15). It seems likely that *C. neoformans* actively manipulates its phagosome to create a permissive intracellular niche; however, to date, no effector proteins directly responsible for such manipulation have been reported. Furthermore, different host immune polarization states have been associated with different infection outcomes (4, 16, 17). M1, or pro-inflammatory, macrophages are associated with clearance or control of infection, whereas M2, or anti-inflammatory, macrophages are associated with progression and dissemination of the disease. Using a ratiometric method that measures the amount of a fluorescent dye released from lysosomes, it was shown that at long time points post-infection (>24 hr) there is increased lysosomal permeabilization in naïve (M0) macrophages relative to activated macrophages, with M1 macrophages showing the least lysosomal damage (14). These were, however, static images taken 24 hr apart from each other, which preclude analysis of the dynamics of this process.

Here we use a highly-specific pH-sensitive dye to label the fungus and follow, in real time, its internalization and maturation of the resulting phagosome. This method allows us to follow the fungus by time-lapse microscopy and directly visualize the pH of its phagosome without the need for wash steps, quencher dyes, or additional sequestered fluorescent molecules. Using this method with *C. neoformans*, a non-filamentous mutant of *C. albicans*, and the non-pathogenic yeast *S. cerevisiae*, we were able to identify and characterize three behaviors: (1) phagosomes that acidify and stay acidic; (2) phagosomes that acidify with subsequent neutralization; and (3) phagosomes that never acidified during the duration of the videos. The last two behaviors were significantly more represented in *C. neoformans* than in the other two fungi. These behaviors represented active manipulation of phagosomal acidification and the kinetics and frequency were influenced by the activation state of the host macrophages. More of the *S. cerevisiae* and *C. albicans*’s phagosomes acidify faster in M1 macrophages relative to M0 and M2. However, that is not seen with *C. neoformans* and, instead, in M2 macrophages less phagosomes acidify and it takes longer. Using a membrane damage reporter, we correlated the abnormal behaviors with phagosomal membrane permeabilization, and we show that this can occur rapidly after internalization. These data demonstrate that *C. neoformans* has the ability to efficiently and rapidly manipulate its phagosome, and offers a direct and rapid way to screen for potential effectors. Given the paucity of strategies used by fungal intracellular pathogens that have been reported, we believe this work will open new ways to investigate this process broadly in fungal pathogens.

## 2 Methods

### 2.1 Strains, cell lines, and growth conditions

Strains used were *C. neoformans* serotype A strain KN99α (18) expressing cytoplasmic mCherry (19), the yeast-locked *C. albicans cph1*/*cph1 efg1*/*efg1* double mutant strain (20), and the *S. cerevisiae* strain BY4741 (21) expressing cytoplasmic mCherry (22). All fungal strains were maintained at -80 °C and grown at 30 °C on yeast-peptone-dextrose (YPD) plates.

The human monocytic cell line THP-1 (ATCC TIB-202) was grown in THP-1 complete media (see Text S1 for recipe) and passed 2 – 3 times per week. Every month a new low-passage stock was thawed and the cells were never used after 12 passages. For differentiation into macrophages, the THP-1 cells were treated as described (23), except that we used 0.16 mM phorbol 12-myristate 12-acetate (PMA, Millipore Sigma).

### 2.2 Labeling of yeast cells with pHrodo Green

Stock solutions of pHrodo IFL Green (P36012, Thermo Fisher Scientific) were prepared in DMSO according to the manufacturer’s instructions and stored at -20 °C in single use aliquots. A fresh aliquot was thawed for each experiment. Fungal cultures grown overnight at 30 °C with shaking (225rpm) were diluted to an OD_600_ of 0.2 and grown for two doublings. The log phase cultures were washed in PBS and opsonized with 40% human serum for 30 min at 37 °C with rotation. Serum was obtained from healthy donors with informed consent under a protocol approved by the University of Notre Dame Institutional Review Board. 10^8^ opsonized fungal cells were resuspended in 100 μL of sodium bicarbonate, and 5 μL of pHrodo Green stock was added to a final concentration of 0.5 mM and incubated in the dark for 30 min at 37 °C with rotation. When using fungi not expressing mCherry, 10 μL of CFW was added to a final concentration of 10 μg/mL and stained at the same time as pHrodo Green. Cells were washed once with PBS and resuspended in imaging media (see Text S1) containing IFNγ (200 ng/mL) or IL-4 (50 ng/mL) when necessary. For a more detailed protocol, see Text S1.

### 2.3 Time Lapse Microscopy

Coincubation of opsonized, pHrodo-stained cells with differentiated THP-1 cells was done in 35mm glass-bottom dishes (MatTek Corporation) as described (22). Briefly, THP-1 cells were seeded at a density of 1.37×10^5^ cells/mL and differentiated with 0.16mM PMA for 48 hr followed by a recovery step of 24 hr without PMA. If required, during the recovery step cytokines were added for polarization of the macrophages. For M1 macrophages, the media contained 200 ng/mL of recombinant human IFNγ (R&D Systems). For M2 macrophages, the media contained 50 ng/mL of recombinant human IL-4 (Shenandoah Biotechnology). Prior to imaging, culture media was replaced with prewarmed imaging media containing appropriate cytokines and opsonized and pHrodo-stained *C. neoformans, C. albicans*, or *S. cerevisiae* at a MOI of 5, 1, or 1, respectively. The dish was immediately transferred to a PeCon (Erbach, Germany) P S compact stage-top incubator preheated to 37 °C with 5% CO_2_. A Zeiss Axio Observer 7, with an Axiocam 506 mono camera and a 40x objective was used to image 2×2 fields of view every 3 minutes for at least 180 minutes.

### 2.4 Flow Cytometry

900,000 THP-1 cells were seeded and PMA-differentiated per well in a 6-well plate. Opsonized and pHrodo-stained *C. neoformans* or *S. cerevisiae* were added as above. An additional T25 flask was seeded and unstained fungi grown to provide negative controls for the flow cytometry analysis. At every timepoint, the wells were washed 3X with DPBS and cells lifted with TrypLE without phenol red (Thermo Fisher Scientific). The cells were collected and resuspended in cold DPBS and immediately analyzed by flow cytometry (BD LSRFortessa).

### 2.5 Immunofluoresence

PMA-differentiated THP-1 cells grown on #1.5 round 12mm glass cover slips (Electron Microscopy Sciences) were challenged with opsonized fungi or treated with cytokines as above. At specified timepoints, the cells were washed with DPBS, fixed in 3.7% formaldehyde in PBS for 10 minutes at room temperature (RT), washed with PBS, and permeabilized with 0.3% saponin in PBS for 10 min at RT. Cells were then washed with PBS and blocked for 60 min in 10% goat serum in PBS. Cells were immunolabeled in 1% goat serum at 4 °C overnight with an antibody against Gal3 (rat; M3/38: sc23938, Santa Cruz Biotechnology), or against CD86 (PA5-88284, Invitrogen) or CD206 (PA5-101657, Invitrogen). Coverslips were then washed with PBS and a corresponding Alexa Fluor-conjugated goat secondary antibody (Invitrogen) was diluted to 1 μg/mL in 1% goat serum in PBS with 100 ng/mL of 4’6-diamidino-2-phenylindole (DAPI; Millipore Sigma) and added to cells for 60 min at RT. After washing with PBS, coverslips were mounted to imaging slides with 5 μL Prolong Diamond Antifade Mountant (Invitrogen) and allowed to cure for at least 24 hours before imaging. Images of fixed cells were acquired using a Zeiss Axio Observer 7, with an Axiocam 506 mono camera and a 100X/1.4 plan-apo oil objective.

## 3 Results

### 3.1 *C. neoformans* exhibits various behaviors with altered phagosomal acidification

To directly visualize pH changes in the fungal containing-phagosome, we used an amine-reactive pH-sensitive dye to decorate the surface of *C. neoformans* or *S. cerevisiae*. This dye, pHrodo Green, has a low pKa, is non-fluorescent at neutral pH but fluorescence rapidly increases as you approach pH of 4, is extremely photostable, and can be used with various platforms such as microscopes, plate readers, or flow cytometers. pHrodo staining does not affect the growth of *C. neoformans* or *S. cerevisiae* (Fig. S1A), under permissive or stress conditions (Fig. S1B), and it is responsive to multiple pH changes without any appreciable loss of fluorescence (Fig. S1C). We incubated serum-opsonized, pHrodo-stained fungal cells with naïve (M0) THP-1 cells, a well characterized human monocytic cell line, and imaged the cells for up to 3 hr under a heated, humidified, stage-top chamber containing 5 % CO_2_ (see Methods). *C. neoformans* and *S. cerevisiae* express soluble mCherry which facilitates the tracking of the cells before and during internalization, but we also used non-fluorescent yeasts stained with calcofluor white (CWF), including a yeast-locked *C. albicans* mutant strain (Fig. S1D). After analyzing *S. cerevisiae* internalization events (n = 60), we found that 76.7% of the events showed rapid acidification that remained for at least 90 minutes (behavior 1). One event (1.6%) lost the acidification before 90 minutes (behavior 2) and the rest (21.7%) never acidified (behavior 3). Stills from the main representative behavior of *S. cerevisiae* are shown in Fig. 1A and the behaviors are quantified in Fig. 2A. In contrast, a substantial number of the *C. neoformans* internalization events showed behavior 2 (16.7%; n = 7) or behavior 3 (26.2%; n = 11), whereas only a bit more than half of the events exhibited behavior 1 (57.1%; n = 24). Examples of these behaviors are shown in Fig. 1B-D and are quantified in Fig. 2B. Notably, the *C. albicans* strain that is unable to form hyphae exhibits the same behaviors as *S. cerevisiae* (Fig. 1G and 2C), suggesting that the alterations to the phagosomal maturation are specific to *C. neoformans*. Consistently, if we use two *C. neoformans* mutant strains known to be defective in intracellular growth, these exhibit mostly behavior 1, just like *S. cerevisiae* or the yeast-locked *C. albicans* (Fig. 1E,F and 2D,E). These findings demonstrate that *C. neoformans* can alter phagosome acidification in various ways in M0 macrophages.

**Fig. 1.**
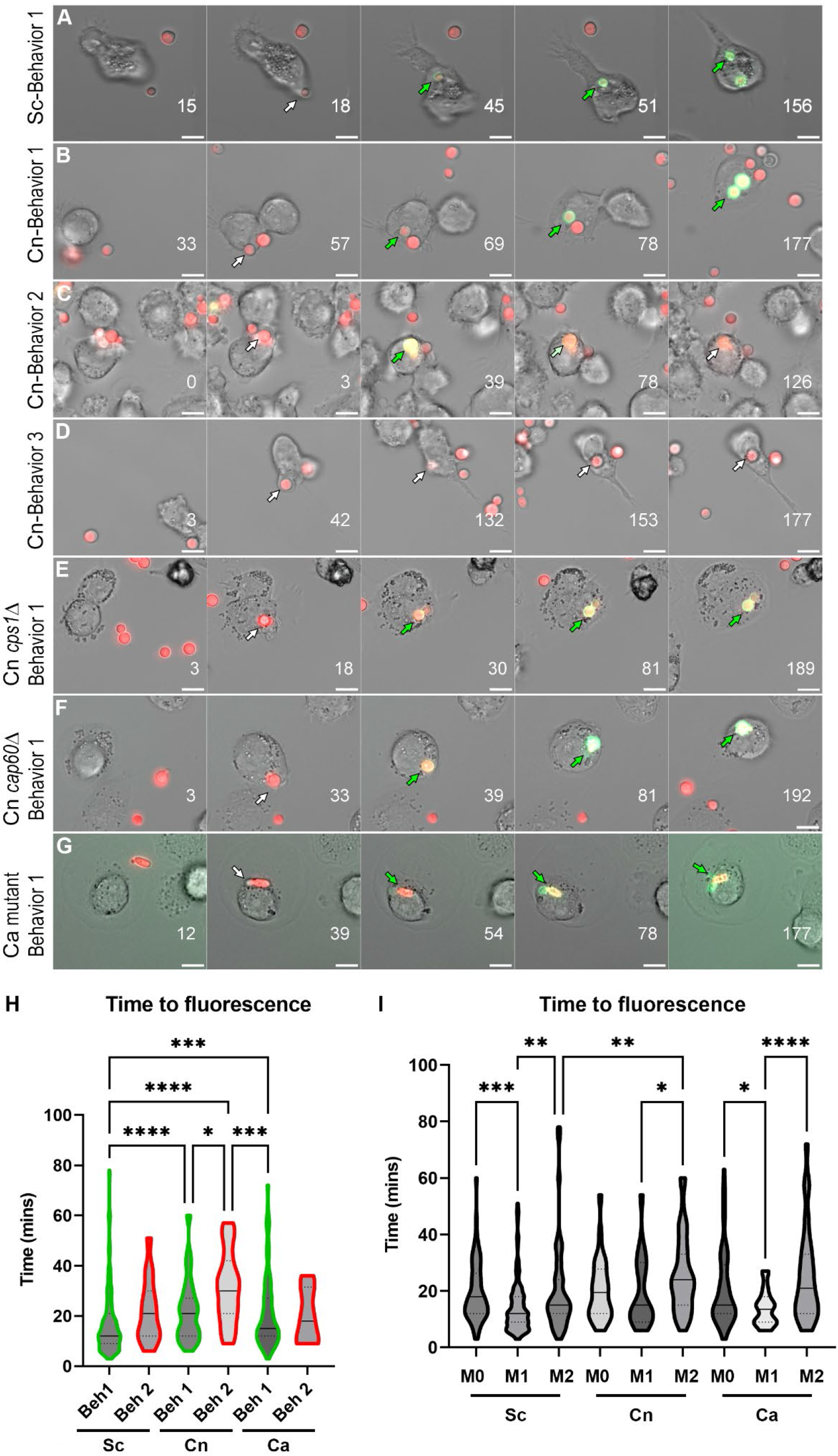
Behaviors and kinetics of phagosomal acidification in fungi-infected THP-1 cells. (A) Stills from a representative video showing the main behavior (behavior 1) in *S. cerevisiae* (Sc)-infected THP-1 cells. For the complete video see Video S1. (B) Stills from a representative video showing behavior 1 in *C. neoformans* (Cn)-infected THP-1 cells. For the complete video see Video S2. (C) Stills from a representative video showing behavior 2 in Cn-infected THP-1 cells. For the complete video see Video S3. (D) Stills from a representative video showing behavior 3 in Cn-infected THP-1 cells. For the complete video see Video S4. (E) Stills from a representative video showing behavior 1 in THP-1 cells infected with Cn mutant *cps1*Δ. For the complete video see Video S5. (F) Stills from a representative video showing behavior 1 in in THP-1 cells infected with Cn mutant *cap60*Δ. For the complete video see Video S6. (G) Stills from a representative video showing behavior 1 in THP-1 cells infected with yeast-locked *C. albicans* (Ca). For the complete video see Video S7. Numbers on lower right represent the time in minutes. The arrows point to the same fungus and are white when nonfluorescent in the pHrodo channel and green when fluorescent. Scale bar = 10 μm. (H) Violin plots showing the kinetics and distribution of behaviors in THP-1 cells infected by Sc, Cn, or Ca. For this analysis, all behavior 1 or behavior 2 events throughout all polarization states were combined. The solid line represents the median, and the dashed lines the quartiles. (I) Violin plots showing the kinetics and distribution of behaviors in THP-1 cells infected by Sc, Cn, or Ca. For this analysis, all behaviors events per polarization state were combined. The solid line represents the median, and the dashed lines the quartiles. See Table S1 for mean and standard deviation values. Violin plots were analyzed by one-way nonparametric ANOVA test, with multiple comparisons (Kruskal-Wallis test). *, P < 0.05; **, P < 0.01; ***, P < 0.001; ****, P < 0.0001.

**Fig. 2.**
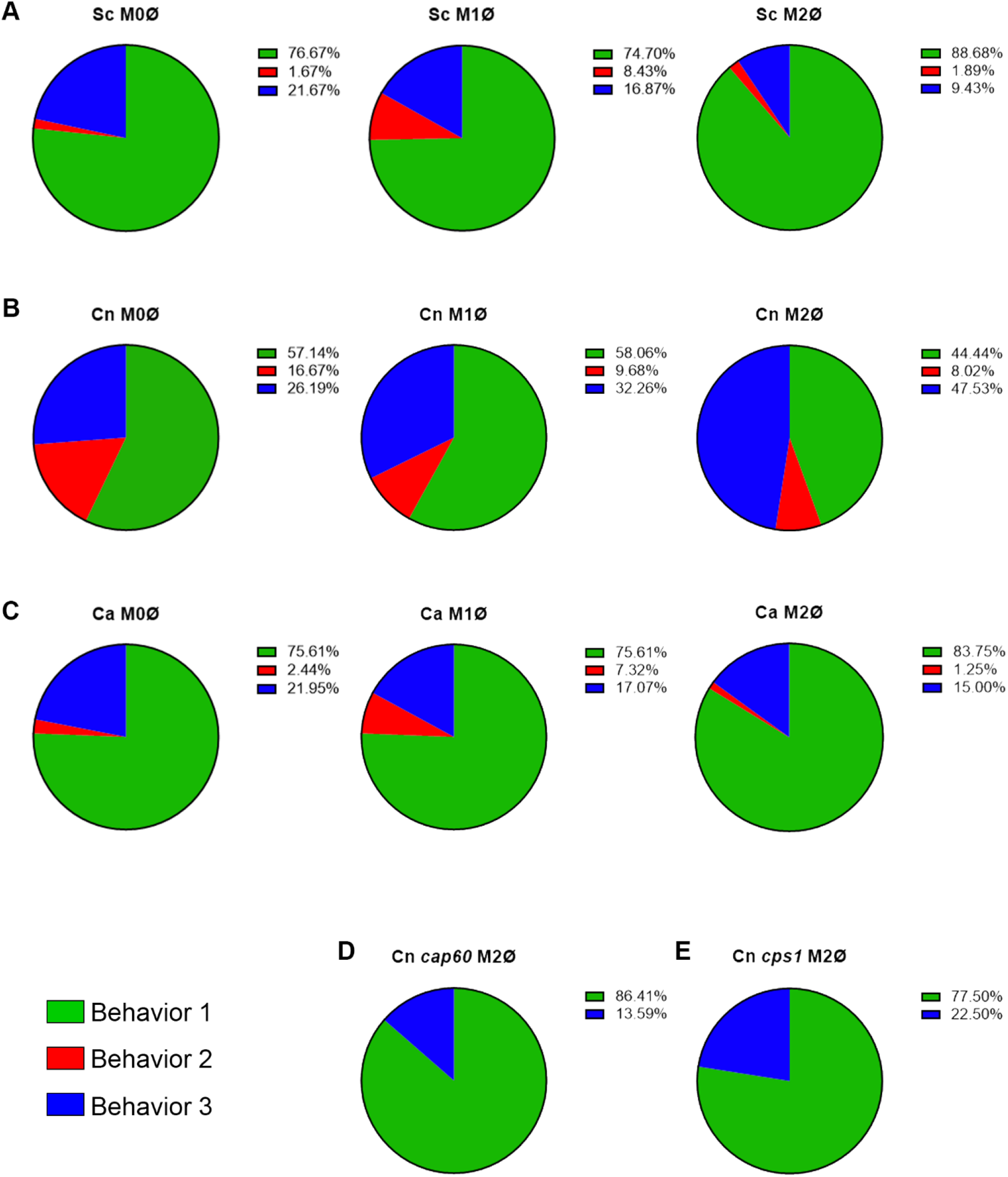
*C. neoformans* actively manipulates phagosomal pH in an immune-dependent way. (A – C) Quantification of all behaviors under all immune conditions of THP-1 cells infected with (A) *S. cerevisiae* (Sc), (B) *C. neoformans* (Cn), and (C) *C. albicans* (Ca). (D and E) Quantification of the behaviors exhibited by Cn mutants (D) *cap60*Δ and (E) *cps1*Δ in M2 macrophages.

### 3.2 The phagosomal acidification behaviors are affected by the polarization state of the host cell

To evaluate if immune signals affected the ability of these fungi to alter acidification of their phagosomes, we repeated the assay with THP-1 cells preactivated to M1 with INFγ, or to M2 with IL-4, using published methods (24). Protein expression of M1 (CD86) and M2 (CD206) markers was confirmed by immunofluorescence (Fig. S2). M1 and M2 macrophages markedly reduced the minor behaviors in *S. cerevisiae* (Fig. 2A; n = 166 for M1 and n = 53 for M2) and yeast-locked *C. albicans* (Fig. 2C; n = 41 for M1 and n = 80 for M2), and still most phagosomes rapidly acidified and stayed acidic. In *C. neoformans*, however, we found that the different polarization states had slightly opposite results (Fig. 2B). In M1 macrophages (n = 31), while most phagosomes still acidify and stay acidic, there was a shift from behavior 2 to behavior 3. In contrast, in M2 macrophages (n = 162), there was a substantial decrease in behavior 1 with a concomitant increase in behavior 3. These findings indicate that *C. neoformans* can manipulate its phagosome in an immune-dependent manner.

### 3.3 Despite similar kinetics, the frequencies and distributions of the *C. neoformans’s* phagosomal acidification behaviors are distinct to the other fungi

To obtain a sense of the kinetics of acidification, we determined how long after phagocytosis the fungi became fluorescent (Table S1). When compared within and between fungi, none of the behaviors’ kinetics were statistically different between the different polarization states. For behavior 1, the overall time to acidification was 16.9± 12.5 min, 22.8± 12.7 min, and 21.3± 14.4 min for *S. cerevisiae, C. neoformans*, and *C. albicans*, respectively; and 22± 12.7 min, 25.3± 13.9 min, and 20± 11.2 min for behavior 2 (Table S1). Likewise, the kinetics are not statistically different between behaviors within host polarization states, however, there was an obvious difference in the pattern of *C. neoformans* compared to the other fungi. For both *S. cerevisiae* and *C. albicans*, M1 polarization had the faster time to acidification, while M0 and M2 had similar kinetics. For *C. neoformans*, on the other hand, M0 and M1 were similar, and M2 had the slowest time to acidification. These results show that the main difference between *C. neoformans* and the other fungi is the distribution of behaviors 2 and 3, and how frequently *C. neoformans* is able to avoid acidification or modulate the pH once its phagosome has acidified, which the other fungi are rarely able to do. To visualize these differences, we generated violin plots of overall times to fluorescence per behavior (Fig. 1H; same behaviors between polarization states combined) or per host immune status (Fig. 1I; all behaviors within the same polarization state combined). *S. cerevisiae*’s behavior 1 was statistically different to the others with 75% of its data points lumped together in one peak in the bottom of the plot (95% CI 12 – 15 min). In contrast, the distribution of *C. neoformans*’s behavior 1 shows 2 peaks, each with about the same amount of data points (95% CI 15 – 24 min). *C. albicans*’s plot also has 1 peak like *S. cerevisiae*, but the data points are spread out more than *S. cerevisiae* (95% CI 15 – 18 min). In terms of host polarization states, for both *S. cerevisiae* and *C. albicans*, M1 resulted in faster acidification (95% CI 11 – 13 min and 9 – 15 min, respectively). In contrast, for *C. neoformans*, M2 resulted in slower acidification (95% CI 21 – 27 min) whereas M1 was not statistically different from M0. This suggests that a significant percentage of the *C. neoformans* population is able to manipulate its phagosome, altering the resulting distribution of phagosomal kinetics.

### 3.4 A significant percentage of *C. neoformans* phagosomes exhibit impaired acidification compared to *S. cerevisiae* phagosomes

To obtain a more general idea of how far reaching *C. neoformans*’s ability to modulate phagosome acidification is, we used the same setup as above but analyzed at least 10,000 events by flow cytometry. The detailed gating scheme is depicted in Fig. S3. This analysis clearly showed the inability of *S. cerevisiae* to avoid phagosome acidification, especially in activated macrophages (Fig. 3A and E). In contrast, a significant percentage of the macrophages infected with *C. neoformans* were unable to acidify their phagosomes (Fig. 3E). This was evident at all timepoints under all polarization states. Notably, an increase in acidified phagosomes over time was seen only in M1 macrophages, while in M0 and M2 the percentage remained relatively constant (Fig. 3E). This is in stark contrast to *S. cerevisiae*, which over time almost 100% of the infected macrophages had acidified phagosomes, despite more of the macrophages being infected than *C. neoformans* (Fig. S3B). This demonstrates a clear ability in *C. neoformans* to modulate phagosome acidification in as short as 30 minutes post-infection, and that this ability is counteracted by host immune signals associated with M1 polarization.

**Fig. 3.**
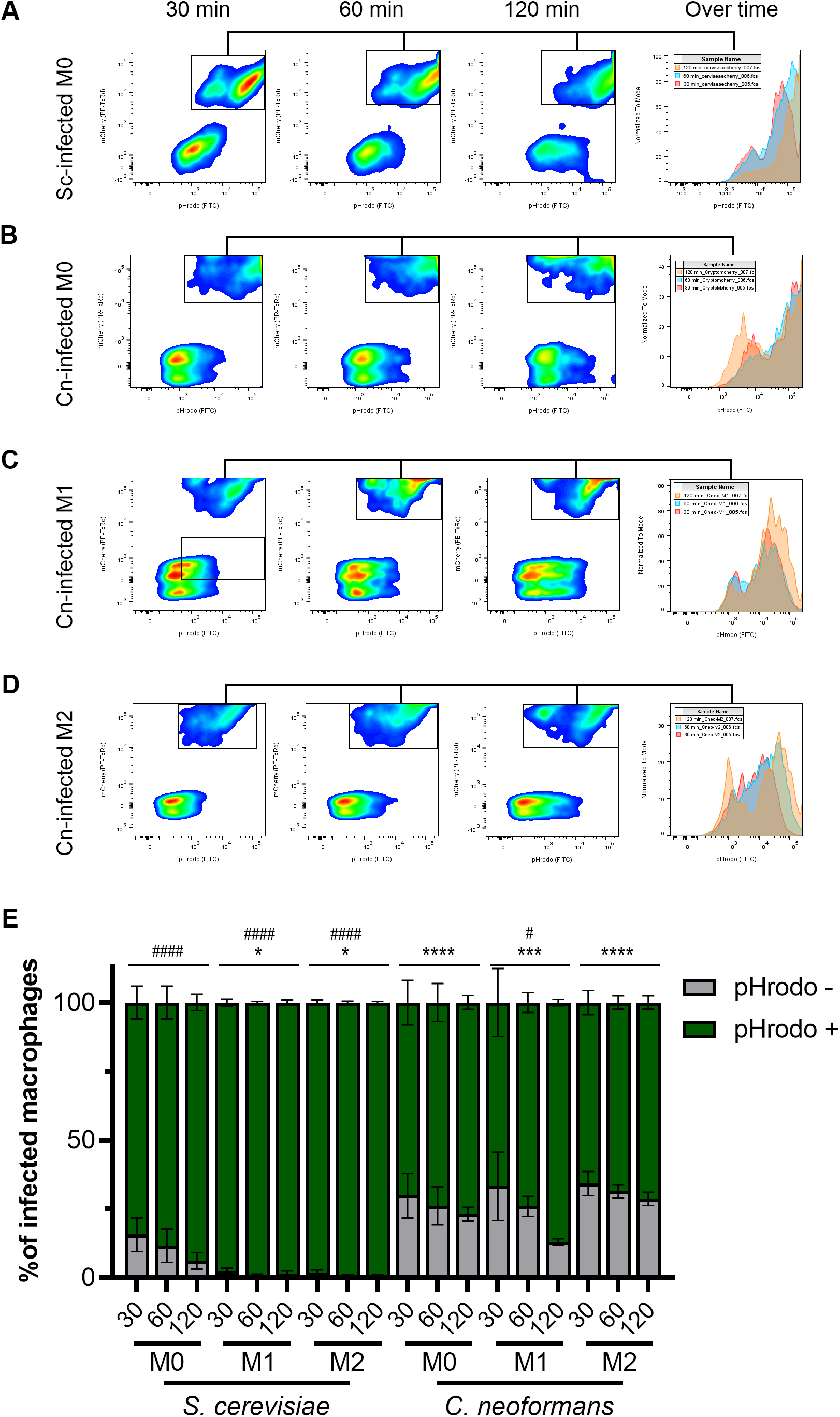
The majority of *C. neoformans* phagosomes exhibit impaired acidification but this is modulated by the macrophage polarization state. (A) Representative flow plots of 30, 60, and 120-minute incubation of *S. cerevisiae* (Sc) with M0 macrophages. The histogram to the right shows the relative number of pHrodo-positive events over time. (B – D) Representative flow plots of 30, 60, and 120-minute incubation of *C. neoformans* (Cn) with (B) M0, (C) M1, and (D) M2 macrophages. The histogram to the right shows the relative number of pHrodo-positive events over time. See Fig. S3 for more details on the gating strategy. Statistics are a 2-way ANOVA with multiple comparisons. The asterisks (*) represent comparisons to *S. cerevisiae*-infected M0 cells. The number sign (#) represent comparisons to *C. neoformans*-infected M0 cells. * or #, P < 0.05; ***, P < 0.001; **** or ####, P < 0.0001.

### 3.5 *C. neoformans* damages and permeabilizes its phagosome early after infection

Previous studies have reported phagosomal permeabilization by indirect means (13-15). They were all looking for the effects of phagosomal permeabilization at long timepoints after infection, usually 24, 48, and 72 hr. We see pH manipulation as fast as 30 min post-infection. One way *C. neoformans* could manipulate phagosome acidification could be by permeabilizing the phagosome membrane, causing neutralization of its contents. We can directly visualize this membrane damage using galectin-3 (Gal-3) staining. Gal-3 will recognize the sugars present inside the phagosome, in the luminal side of its membrane, but only if the membrane is damaged (25, 26). Hence, when the phagosome is permeabilized, Gal-3 has access to its contents and will decorate the interior side of the phagosome (Fig. 4A). In this way, Gal-3 acts as a direct membrane damage reporter. We did a timecourse with *C. neoformans*-infected M0 macrophages and saw little colocalization with the cryptococcal-phagosome at 1 hr, but that increased over time, peaking at 8 hr post-infection (Fig. 4B). We chose this timepoint to test if host immune status also affected this membrane damage similar to how it affected pH modulation by the fungus. We found that about one-third of the cryptococcal-phagosomes were decorated with Gal-3 in M0, and that dropped to 17.9% in M1, whereas in M2 it increased to 50% (Fig. 4C). In comparison, when infected with *S. cerevisiae*, only 10% (M0), 7% (M1), or 18% (M2) of the phagosomes are positive for Gal-3. The incidence of membrane damage found with Gal-3 staining correlates well with the percentage of pHrodo-negative macrophages found by flow cytometry or combined behaviors 2 and 3 found by live microscopy. This suggests that *C. neoformans*’s ability to damage its phagosome might be one of the strategies used by this fungus and it happens early on cellular infection.

**Fig. 4.**
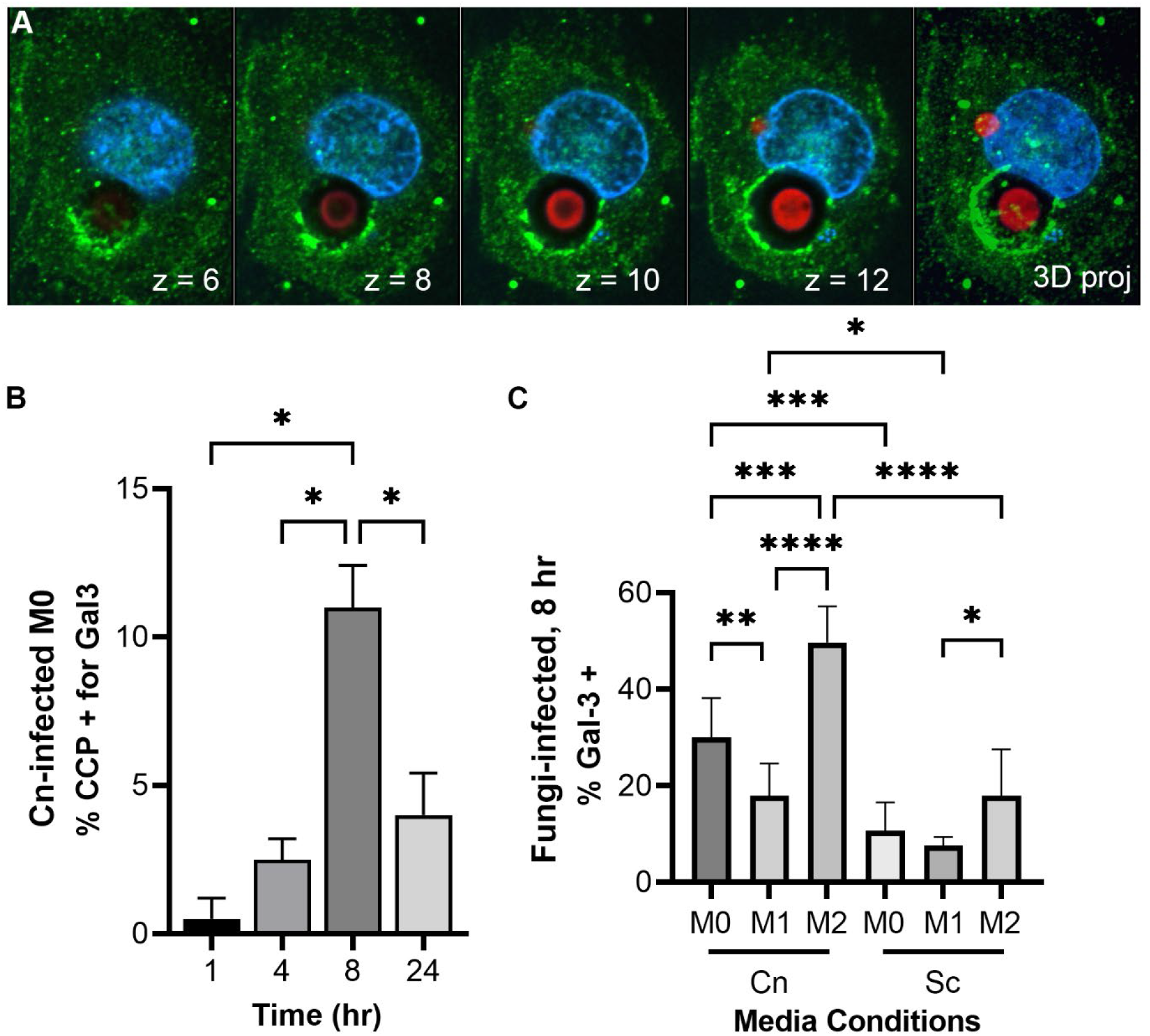
*C. neoformans* damages and permeabilizes its phagosome early after infection in an immune signal-dependent way. (A) Images from a representative THP-1 cell (nuclei in blue) 24 hr-post infection with *C. neoformans* (red) showing membrane damage around one cryptococcal-phagosome (Gal-3 in green). The number on the lower right represent the Z-slice on a series of pictures that are reconstructed into a full projection on the last image to the right. The dark halo surrounding the cell represents its polysaccharide capsule. (B) Quantification of Gal-3 staining over time in M0 THP-1 cells infected with *C. neoformans* (Cn). CCP, cryptococcal-containing phagosome. (C) Quantification of Gal-3 staining at 8 hr in M0, M1, and M2 THP-1 cells infected with Cn or *S. cerevisiae* (Sc). Statistics are ordinary one-way ANOVA with multiple comparisons. *, P < 0.05; **, P< 0.01; ***, P < 0.001; ****, P < 0.0001.

## 4 Discussion

Here we use a novel pH-sensitive staining procedure to follow and directly visualize changes in pH in fungal-containing phagosomes. We applied this method to fluorescent strains of *C. neoformans* and *S. cerevisiae*, and analyzed them by live microscopy, allowing us to identify different behaviors and measure their dynamics, as well as by flow cytometry, which allows us to capture and analyze tens of thousands of events. To demonstrate broader applicability of the method, we also used non-fluorescent strains of *C. neoformans* and *C. albicans*, highlighting the possibility of using this dye to study other intracellular fungal pathogens. This dye and similar approaches have recently been used to study phagocytosis, localization, and effects of bacterial infections on host cells (27-30). Although it is likely that *C. neoformans* somehow alters its phagosome, in regards to acidification, published reports are controversial. Moreover, to date no cryptococcal effector necessary for this phagosomal manipulation has been identified. Here we show that a significant percentage of the cryptococcal phagosomes do not acidify normally, and this alteration is dependent on the immune status of the host phagocyte. This is not surprising given that different polarization states have different effects on inflammation (31). However, all macrophages, despite polarization, acidify their phagolysosomes to the same extent, albeit with different kinetics (32). This is seen, for example, with *S. cerevisiae* and the yeast-locked *C. albicans*, that regardless the polarization state, most of their phagosomes acidify. In contrast, with *C. neoformans*, a majority of the phagosomes do not acidify, and that observation is exacerbated in M2 macrophages.

Furthermore, for the first time, we directly visualize, and quantify, phagosomal membrane damage, which might be one of the strategies used by *C. neoformans* to alter phagosomal acidification, but it is not the only one. Phagosomal permeabilization is a common mechanism used by intracellular bacterial pathogens, where effectors that cause membrane damage are essential for their intracellular survival. Gal-3 staining has been used to show that *Shigella* and *Salmonella* damage their phagosomes in order to escape into the cytoplasm (26, 33). *C. neoformans*, however, remains inside a phagosome, but damaging the membrane to access the host cytoplasm would represent a big advantage. Nevertheless, phagosomal membrane damage does not seems to be completely essential as in the first 2 – 3 hr post-infection, up to 34% of the phagosomes are not acidic (by flow cytometry; Fig. 3E) and up to 55% of the phagosomes exhibit some alteration (by live microscopy; Fig. 2B), yet only ∼5% of cryptococcal-phagosomes stain with Gal-3 in that timeframe. Clearly, there are other mechanisms used by this fungus to manipulate phagosomal acidification. Nevertheless, both of these phenotypes (large percentage of pHrodo-negative phagosomes and phagosomes decorated with Gal-3) are mostly absent from *S. cerevisiae*, a fungus of similar morphology and size, demonstrating that *C. neoformans* actively cause these alterations. The fact that there are multiple behaviors, including behavior 1 which is shared with *S. cerevisiae*, may explain the different results that have been reported about the cryptococcal phagosome (Fig. S4).

Interestingly, *C. neoformans* is known to escape phagocytes by non-lytic exocytosis (NLE), a phenomenon whereby the fungal cells are secreted from the host cell without any adverse effect to any cell type (34-36). This has been reported to happen in about 10 – 30% of the infected cells, which is similar to the percentage of infected macrophages that are pHrodo negative in our flow cytometry studies (Fig. 3E) and the percentage of phagosomes that are decorated with Gal-3 (Fig. 4C). Phagosomal acidification has been shown to affect NLE, with NLE events happening from nonacidified phagosomes (9). Notably, we captured two NLE events where the pHrodo signal was lost and shortly thereafter the fungi underwent NLE (Video S8). The fact that we only saw 2 events in our live imaging is not surprising since the Casadevall group has shown that these events usually happen after several hours of intracellular residence (>6 hr) while we only imaged for up to 3 hr (37). Nevertheless, our findings are in line and consistent with NLE reports, providing additional support that NLE is an active process triggered by *C. neoformans*’s manipulation of its phagosome.

M1 and M2 macrophages exhibit different abilities controlling cryptococcal intracellular growth; however, it is not known if these differing outcomes correlate with different fungal intracellular niches. Interestingly, we show that the frequency of phagosome manipulation by *C. neoformans* varies between the different host activation states. Although in our flow cytometry analysis only the M1 macrophages were statistically different to other conditions, we could clearly see a trend in the M2 macrophages where the population of pHrodo negative phagosomes was higher than in M0. We acknowledge that the analysis by flow cytometry is limited in the sense that not all phagosomes within the same host cell are manipulated equally; however, the fact that with *S. cerevisiae* the pHrodo-positive population increased to almost 100% in all conditions whereas with *C. neoformans* this population remains constant argues that the trend seen in M2 macrophages is relevant. Consistently, when analyzing the kinetics of acidification in the different polarization states, M2 had the higher acidification times relative to M0 or M1 (Fig. 1I). Moreover, the differences in manipulation over time in M1 relative to M2 macrophages correlate with animal studies where M1 macrophages have been shown to control infection better than M2 macrophages. Hence, our studies support the hypothesis that the ability to manipulate phagosomal maturation correlates with the infection outcome. Consistently, cryptococcal *cap60*Δ and *cps1*Δ mutants, which cannot grow intracellularly and are avirulent, cannot alter acidification of their phagosomes (Fig. 2D, E).

In conclusion, here we demonstrate that *C. neoformans* can manipulate acidification of its phagosomes in various ways, that this ability is affected by the immune status of the host, and that one way this manipulation could happen is by phagosomal membrane permeabilization. We present a versatile and easy method to screen phagosomal changes by live microscopy or by flow cytometry. We show that *C. neoformans*’s ability to manipulate its phagosome is robust and happens soon after internalization. Analysis of cryptococcal mutants with these methods might identify effectors necessary for this manipulation, but these methods can be broadly applicable to other intracellular yeasts as well.

## 5 Author Contributions

FHS-T was responsible for the study conception and design. EJS-B, PVS, and FHS-T performed the experiments, collected data, and analysed results. PVS and FHS-T prepared the figures and wrote the manuscript with input from EJS-B. All authors read the manuscript and provided edits and feedback. EJS-B and PVS share first authorship and the naming order was determined alphabetically.

## 6 Funding

This work was supported by start-up funds from the University of Notre Dame to FHS-T. EJS-B was partially supported by a Kinesis-Fernández Richards Family Fellowship.

## 7 Acknowledgments

We thank members of the Santiago-Tirado lab for comments and feedback on this paper.

## 9 Figure Legends

**Fig. S1.**
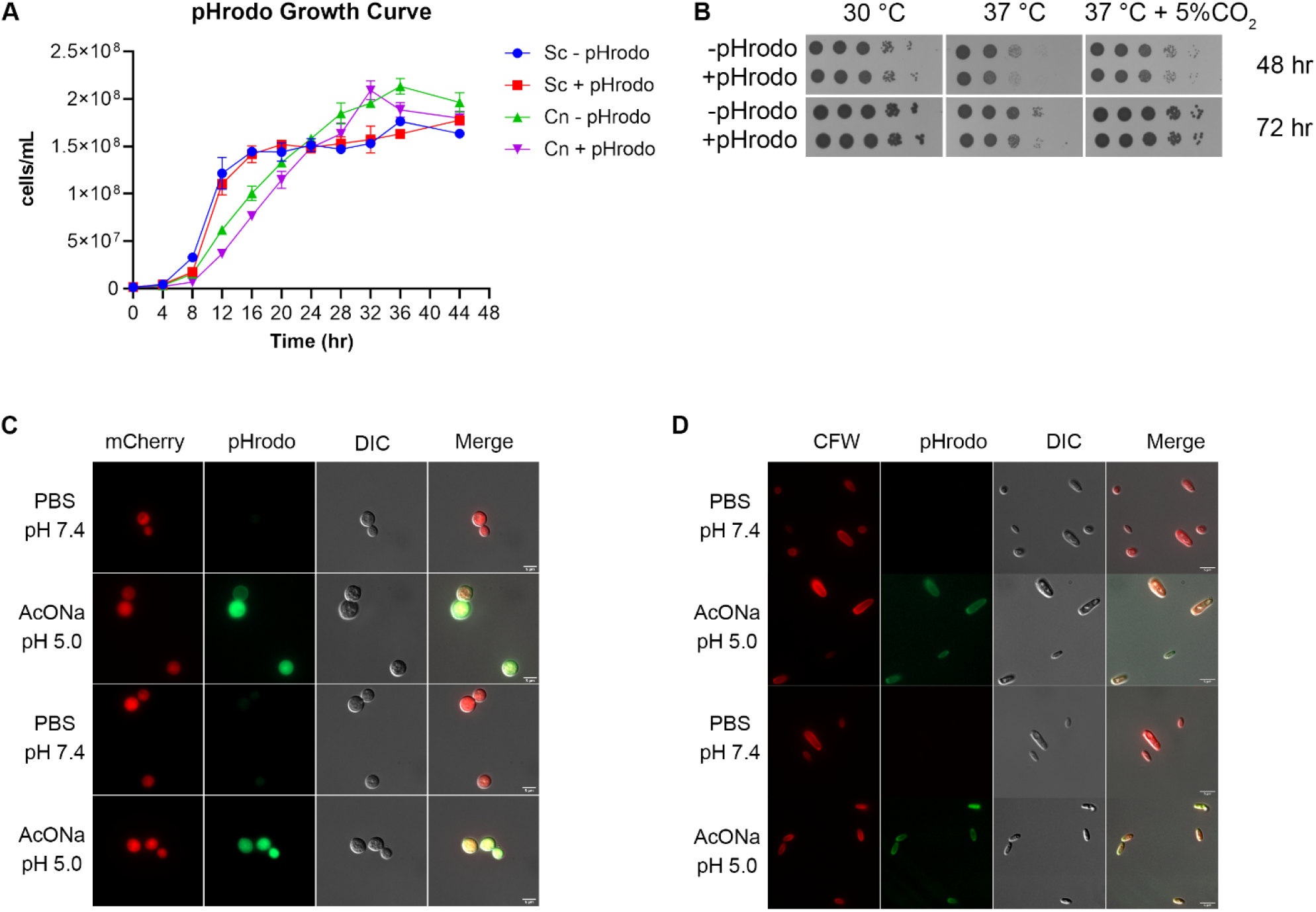
Characteristics of pHrodo-stained fungal cells. (A) Growth curve of unstained and pHrodo-stained *S. cerevisiae* (Sc) and *C. neoformans* (Cn) in YPD media at 30 °C over 48 hr. No effect of pHrodo staining is seen in either of the fungi. (B) Dot spot growth analysis in RPMI-agar plates of Cn under the indicated conditions. Images were taken at 48 and 72 hr. (C and D) Representative images of pHrodo-stained (C) Cn and (D) yeast-locked *C. albicans* (Ca) cells after sequential resuspension in PBS, pH 7.4, and sodium acetate buffer (AcONa), pH 5.0, showing that the pHrodo staining is responsive to changes in pH. Ca cells are not fluorescent and were counterstained with CFW. The CFW fluorescence was artificially colored red for ease of view. Scale bars represents 5 μm.

**Fig. S2.**
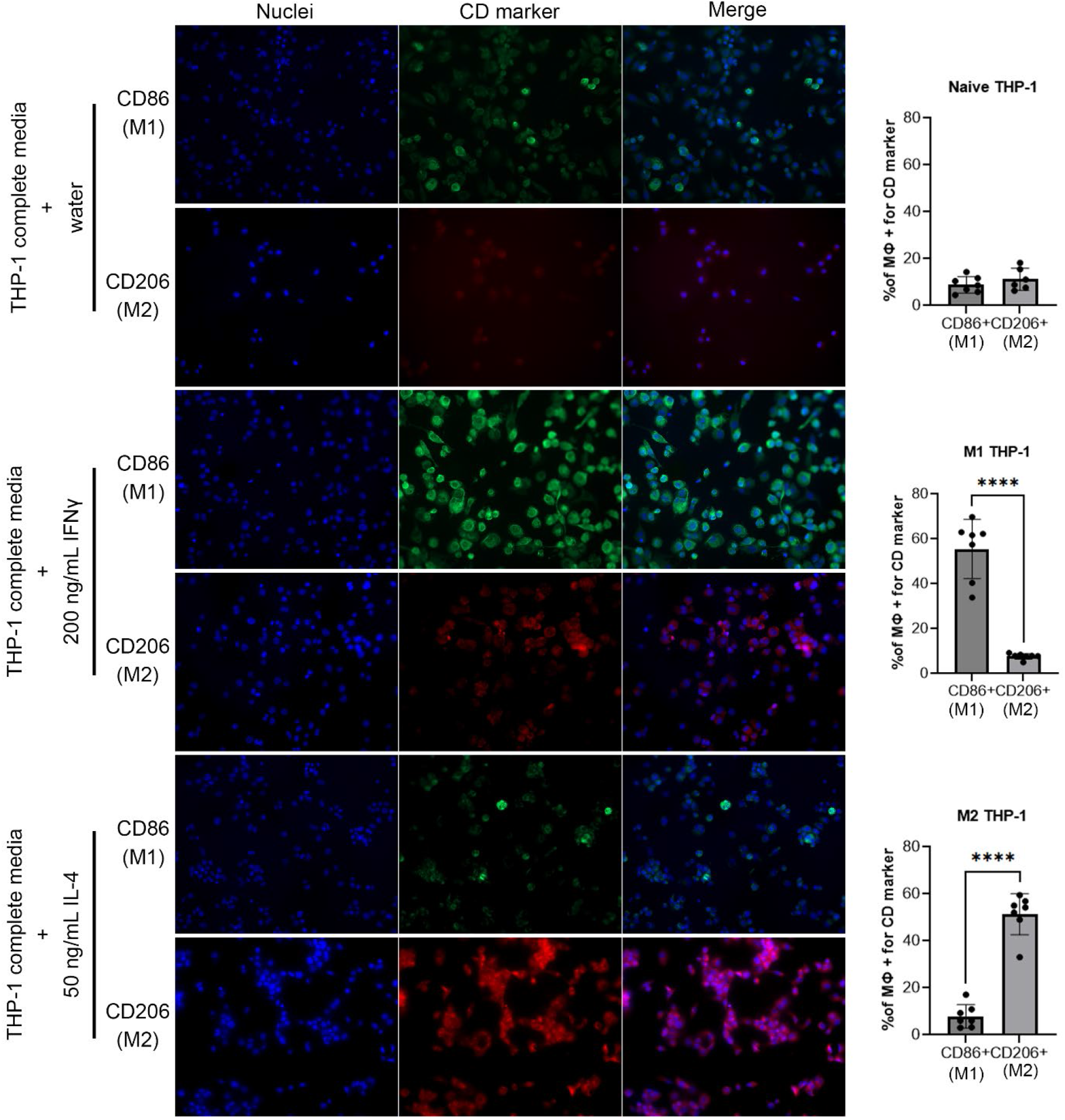
Polarization of THP-1 cells using IFNγ or IL-4 treatment. THP-1 cells were treated with IFNγ, IL-4, or vehicle (water) as described in the Methods section. Shown are representative fields of views of immunofluorescence analysis using DAPI (blue) to stain nuclei; anti-CD86 antibody (artificially colored green); and anti-CD206 antibody (artificially colored red). The percentage of cells positive for each marker is quantified on the right. Each circle represents a coverslip, 2 coverslips per biological independent experiment. Statistics are unpaired t-tests comparing the two conditions. ****, P < 0.0001.

**Fig. S3.**
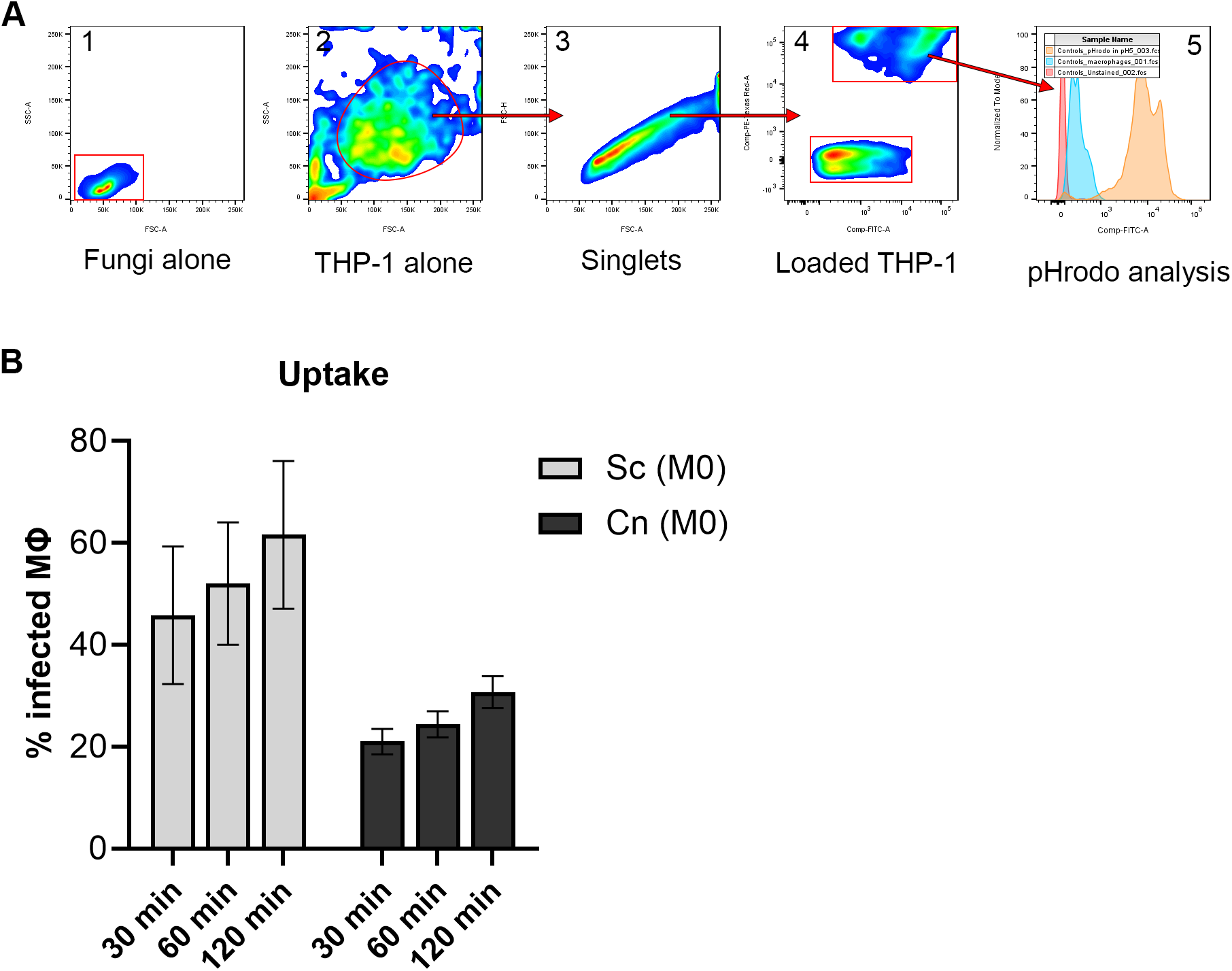
Flow cytometry analysis. (A) Representative plots showing the flow cytometry gating strategy. First, unstained fungal and host cells are analyzed to determine their position in the size-scatter plot. These positions are used to eliminate free fungi (and cell debris) and select healthy THP-1 cells (gates 1 and 2). The healthy THP-1 cell population is used to select for single cells (gate 3). The singlets are then used to select for the population positive in mCherry (gate 4). The population negative to mCherry are uninfected THP-1 cells. Lastly, the infected population is used to analyze pHrodo signal (gate 5). (B) Uptake of fungal cells from a representative flow experiment using M0 macrophages. Despite using only an MOI of 1 (versus an MOI of 5 for *C. neoformans*), *S. cerevisiae* (Sc) is phagocytosed more avidly than *C. neoformans* (Cn). This data comes from Gate 4, the mCherry-positive population represents infected macrophages. The values in here were calculated by dividing gate 4 by gate 3.

**Fig. S4.**
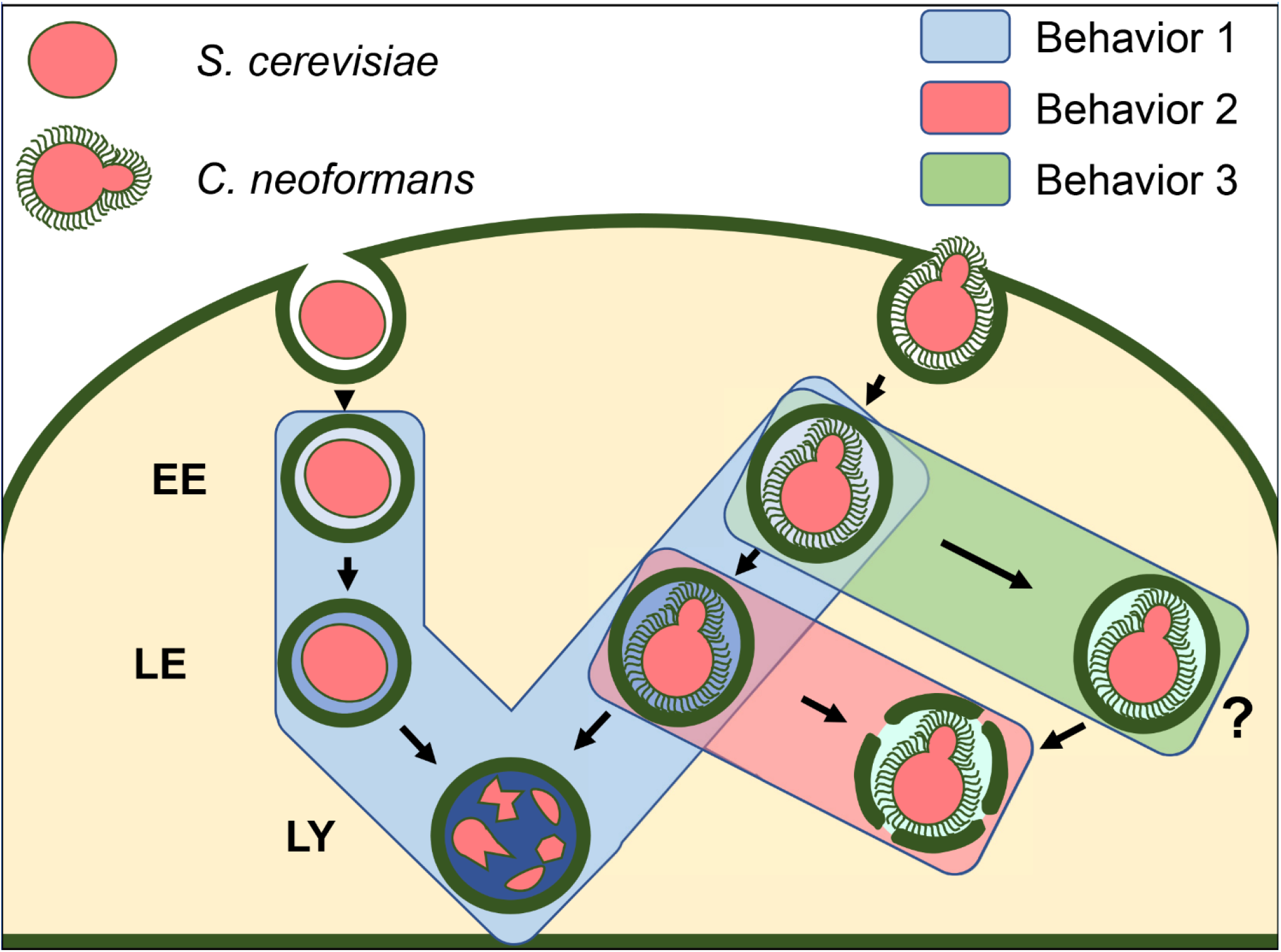
Model summarizing our findings. We found three behaviors in cells infected with either *S. cerevisiae* (unbudded pink cell) or *C. neoformans* (budded pink cell surrounded by capsule). Behavior 1 (blue path) is shared between the two fungi and presumable results in formation of a fully-functional phagolysosome, where the fungal cells are destroyed. Behavior 2 (red path) deviates from the normal path and results in loss of acidification. This can happen in part by phagosomal membrane permeabilization (broken dark green outline). Behavior 3 (green path) results in cells that never acidify, hence this path represents an unknown phagosomal compartment (depicted by ‘?’). This compartment never acidifies so it could also result in phagosomal permeabilization but we would not be able to see it in our live imaging.

**Table S1**. Kinetics of acidification of fungal phagosomes (in minutes).

**Text S1**. Supplementary Methods.

**Video S1**. Representative video of the main behavior (behavior 1) of *S. cerevisiae* phagosomes.

**Video S2**. Representative video of behavior 1 of *C. neoformans* phagosomes. **Video S3**. Representative video of behavior 2 of *C. neoformans* phagosomes. **Video S4**. Representative video of behavior 3 of *C. neoformans* phagosomes.

**Video S5**. Representative video of behavior 1 of *C. neoformans* mutant *cps1*Δ. This mutant was stained with CFW, but the color was changed to red for ease of visualization.

**Video S6**. Representative video of behavior 1 of *C. neoformans* mutant *cap60*Δ. This mutant was stained with CFW, but the color was changed to red for ease of visualization.

**Video S7**. Representative video of behavior 1 of yeast-locked *C. albicans* mutant. This mutant was stained with CFW, but the color was changed to red for ease of visualization. Interestingly, the pHrodo dye (green) seems to be inherited by the growing bud, whereas CFW (red) is retained in the mother cell.

**Video S8**. Video depicting two instances of non-lytic exocytosis (NLE). Notice that immediately after losing fluorescence (pH neutralization) the yeast cells are expelled from the host.

## Notes

### Competing Interest Statement

The authors have declared no competing interest.

